# Multiplexed live visualization of cell fate dynamics in hPSCs at single-cell resolution

**DOI:** 10.1101/2021.01.30.428961

**Authors:** Sungmin Kim, Edward Ren, Paola Marco Casanova, Eugenia Piddini, Rafael Carazo Salas

## Abstract

Live imaging can provide powerful insights into developmental and cellular processes but availability of multiplexable reporters has been limiting. Here we describe ORACLE, a cell fate reporter class in which fluorescent proteins fused with the nucleoporin POM121 are driven by promoters of transcription factors of interest. ORACLE’s nuclear rim localisation therefore enables multiplexing with conventional nuclear reporters. We applied ORACLE to investigate the dynamics of pluripotency exit at single-cell level, using human pluripotent stem cells (hPSCs) imaged by multi-day time-lapse high-content microscopy. Using an ORACLE-OCT4 pluripotency marker we reveal that G1 phase length and OCT4 level are strongly coupled and that spatial location in a colony impacts the timing of pluripotency exit. Combining ORACLE-OCT4 and an ORACLE-SOX1 early neuronal differentiation marker, we visualize in real-time the dynamics of cell fate transition between pluripotency and early neural fate, and show that pluripotency exit and differentiation onset are likely not tightly coupled in single-cells. Thus ORACLE is a powerful tool to enable quantitative studies of spatiotemporal cell fate control.

## INTRODUCTION

Human pluripotent stem cell (hPSC) technologies stand to enable the regenerative medicine of the future ^1^. Although hPSCs can in principle be differentiated into almost all cell types in our bodies ^2–5^, targeted differentiation often also leads to generation of undesired cell types and cellular heterogeneity incompatible with safe and effective cell therapies ^6^. Understanding the origin of such heterogeneity is critical to learn how to efficiently, specifically, and safely produce hPSC-derived cells and tissues for both basic and therapeutic applications.

Single-cell RNA sequencing (scRNAseq) studies of hPSCs differentiating into many different cell types - such as retina, cardiac or endodermal cells - have revealed how complex and poorly understood this heterogeneity really is ^7–10^. Although lineage trajectories can be inferred computationally, scRNAseq approaches are terminal and therefore unable to access some fundamental aspects of the spatiotemporal dynamics of cell fate control. These include the dynamics of transcription factor (TF) heterogeneity ^11^ in cells, and the real-time sequences, durations, and spatial changes in TF levels through tissues, as well as key cell biological aspects of fate dynamics like cell movement, cell death and progression through the cell cycle and how they are coordinated and linked with fate. Such dynamical and cell biological layers are only accessible by ‘live’ microscopy ^12,13^. Hence, refined live high-content microscopy tools and approaches are needed to extract this critical information in order to clarify the mechanisms controlling cell fate establishment and how to harness those mechanisms to optimize tissue derivation.

An example illustrating this need is the cell cycle, which is known to regulate fate control in hPSCs. It has been shown that CyclinD/CDK4-6 controls hPSCs differentiation capacity during the G1 phase ^14,15^, while, CyclinB1 promotes pluripotency maintenance in G2, at least for human embryonic stem cells (hESCs) ^16^. Reciprocally, pluripotency factors have also been shown to provide feedback to cell cycle control. For instance, the pluripotency determinants OCT4 and SOX2 control miR-302 expression - a CyclinD1 translational repressor - in G1 ^17^, and OCT4 down-regulation blocks cell cycle progression in G0/G1 via p21 ^18^. These findings derived from cell population studies provide evidence that hPSCs coordinate cell cycle entry with early cell fate changes, yet thus far it has not been possible to investigate this coordination directly. Such studies require tools allowing visualization of both cell cycle and cell fate status at a single-cell level in real-time.

In the case of cell fate dynamics, it has been shown that TF heterogeneity can greatly impact early cell fate decisions during development ^19^ as well as clinical tissue design^20,21^. Recently pioneering studies have begun to explore temporal TF dynamics and cell-to-cell TF heterogeneity in live single cells using transgenic reporter cell lines and quantitative timelapse microscopy ^22^. For instance, continuous long-term cell lineage tracking of mouse embryonic stem cells (mESCs) expressing VENUS-tagged pluripotency TFs Oct4 or Nanog allowed quantification and comparison of their protein expression changes. This analysis revealed a strong heterogeneity in Nanog expression between cells and mESCs subsets with differing Nanog levels, with both factors differently impacting TF correlation networks and differentiation propensities ^23^. Similarly, quantitative multi-generational analysis of an OCT4-mCherry fusion in hESCs revealed that cells’ pre-existing OCT4 pattern can predict their extraembryonic mesoderm differentiation propensity ^24^. Additionally, in a more recent study, quantitative live monitoring of mESCs co-expressing the two pluripotency TF fusion proteins Sox2-SNAP and Oct4-Halo showed that small endogenous fluctuations of Sox2 and Oct4 with respect to each other can bias early neuroectodermal versus mesendodermal fate decisions ^25^. This illustrates the power of integrating live dynamical information from multiple TFs in clarifying otherwise inaccessible layers of cell fate control.

At a general level, analyses that resolve multiple spatiotemporally integrated mechanisms live remain rare, in part because live multiplexable cell reporters compatible with simultaneous visualization are scarce. This is particularly limiting for TF analysis, as TF reporters are all localised to the nucleus, which means that typically few different factors (labelled by fluorophores that emit in different colours) can be co-visualized and distinguished within cells. In addition, functionality/physiology can be impacted when TFs are tagged ^22^. Furthermore, frequent time-lapse, multi-day imaging - as required to track singlecells and quantitatively monitor their cell state and fate changes across generations - can be particularly phototoxic and damaging to delicate cells such as stem cells ^13,26,27^. These issues combined have made it particularly difficult to apply such approaches to hPSCs, and to our knowledge, no study to date has visualized live the spatiotemporal dynamics of multiple TF in human stem cells at single-cell resolution.

Here, we develop a novel class of ‘live’ reporters of TF status named ORACLE, which are localised to the nuclear envelope, making them multiplexable and compatible with conventional nuclearly-localised TF or cell proliferation reporters. We demonstrate their power and broad applicability using classical experimental paradigms in stem cell biology: exit from the pluripotent state and onset of early neuroectodermal differentiation in hPSCs. By establishing hPSC lines expressing ORACLE reporters alone or in combination with conventional TF or cell proliferation reporters, we are able to reveal the spatiotemporal dynamics of cell fate transitions in hPSCs over multiple days and at single-cell level, by optimised multi-day/multi-colour time-lapse high-content microscopy. Using an ORACLE-OCT4 reporter, we show that there is significant cell-to-cell heterogeneity in the temporal loss of pluripotency among hPSCs differentiating into early neural lineage, and that spatial location likely determines the timing of pluripotency exit. By co-visualizing ORACLE-OCT4 and the nuclear cell cycle reporter FUCCI, we demonstrate that cell cycle duration is a poor predictor of pluripotency exit and that, instead, G1 phase length and OCT4 levels are coupled during pluripotency exit, suggesting that they are mechanistically linked. Finally, by combining ORACLE-OCT4 and an ORACLE-SOX1 reporter, we visualize in real-time the transition between pluripotency and early neural differentiation onset and reveal that pluripotency exit and differentiation onset are not tightly coupled in single-cells.

## RESULTS

### ORACLE cell fate reporter design

In order to expand the available toolkit for cell fate monitoring, we set out to establish a new class of fluorescent cell fate reporters. We wanted these to be amenable to multiplexing and visually compatible with commonly used existing reporters of spatiotemporal cell dynamics. All existing reporters - such as fluorescently-labelled histone H2B chromatin ^28^, the FUCCI cell cycle reporter ^29^ and fluorescently-labelled transcription factors (TFs) - are restricted to the nucleus. We therefore reasoned that fate reporters localised to other subcellular localisations would be valuable, complementary tools to enable cell biological discoveries. We considered designing fate reporters localised to cellular compartments such as microtubules, the Golgi apparatus, or the plasma membrane, but reasoned that robustly assigning such reporter signals to specific cells growing in compact colonies, such as hPSCs, would be technically challenging. We instead focused on the nuclear rim, as signals from reporters localised there would likely be clearly distinguishable from nuclear reporter signals and easy to assign to the cell of origin. We trialled several fluorescent protein fusions including KASH domains and LEM proteins ^30,31^, nuclear lamina proteins (Emerin, Lamin A and B ^32,33^) and nuclear pore proteins ^34^, and eventually selected the nuclear pore protein POM121. Relative to the other candidates, POM121 has ideal nuclear-rim protein properties: (a) fluorescently tagged expression in cells does not lead to any detectable morphological defects ^34^, (b) it has well characterized and quality controlled published fusions ^35^, (c) it has little to no diffuse cytoplasmic or nuclear signal, allowing for a distinct nuclear rim localisation, and (d) it has a high enough protein turnover ^34,36,37^ that it could be used as a proxy to monitor transcription factor turnover ^24^.

Based on these properties, we assessed the suitability of fluorescently-labelled POM121 as a reporter of transcription factor level in hPSCs (Fig. 1a and b, left, and Supplementary Fig. 1a and b). In our first strategy (Fig. 1a, left), we generated “exogenous” expression constructs in which fluorescently-labelled POM121 expression can be driven from the TF promoter of choice, by integrating a cassette comprising the TF promoter sequence and the labelled POM121 sequence at genomic safe harbours such as AAVS1 and ROSA26, using CRISPR knock-in (see Methods for detailed descriptions from here on). This approach allows the original TF to function with minimal disruption, provided a promoter sequence is known and well defined for the TF of interest. In the second strategy (Fig. 1b, left), we generated “endogenous” constructs allowing fluorescently-labelled POM121 expression to be controlled directly by the endogenous TF’s promoter, by integrating the labelled POM121 sequence directly at the TF locus via translational coupling using the 2A system ^38,39^. This ensures that expression of the endogenous TF is not affected. We call these types of constructs ORACLE for their ability to reveal fate.

**Figure 1.**
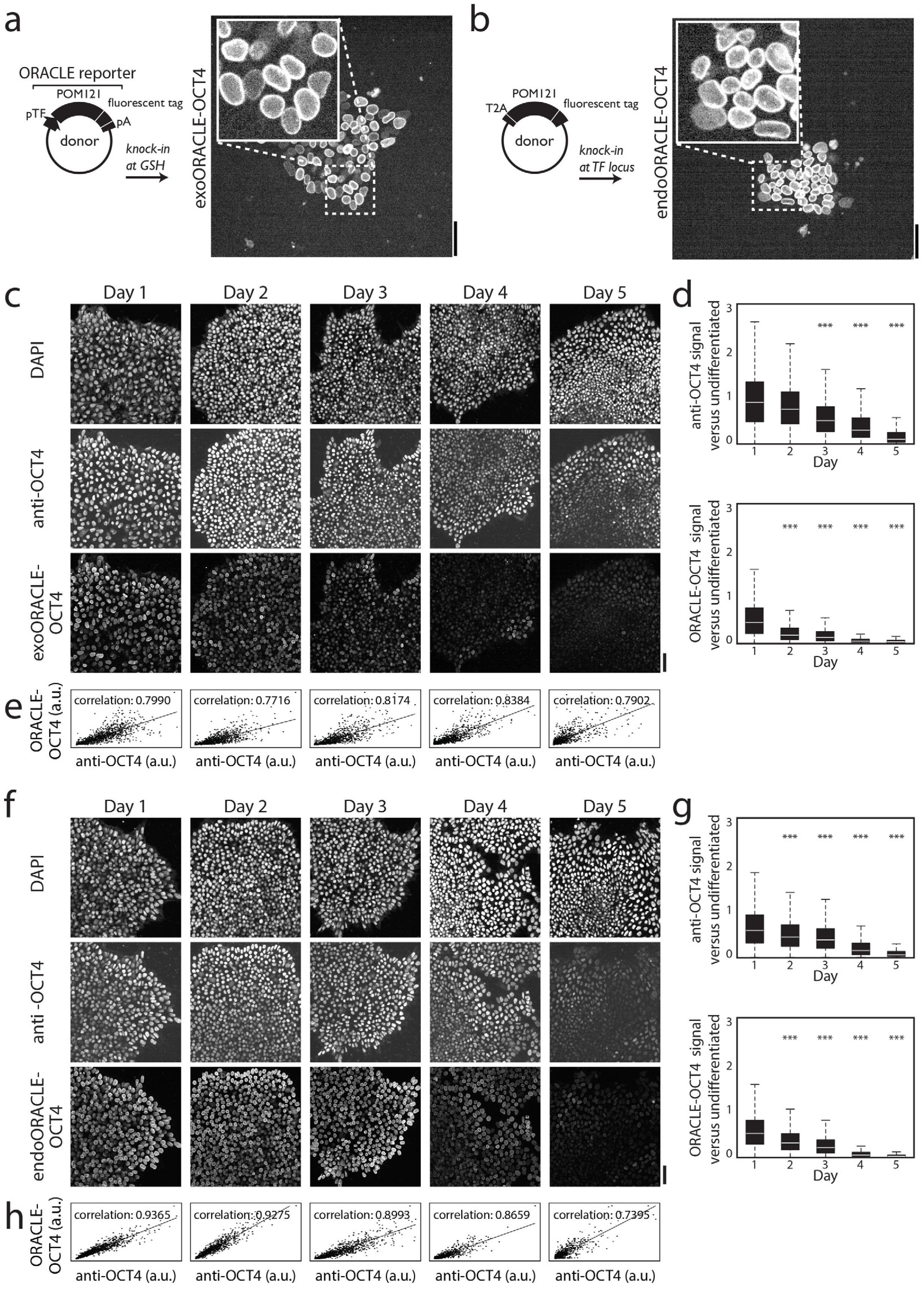
ORACLE-OCT4 level and loss mirror those of OCT4 protein in differentiating hPSCs. **a** and **b,** Left: Schematic of exoORACLE (**a**) and endoORACLE (**b**) reporter design. Right: images of hPSCs stably knocked-in with exoORACLE-OCT4 (**a**) or endoORACLE-OCT4 (**b**). Scalebars: 50 μm. **c, d, f** and **g**, ORACLE-OCT4 level in cells visually (**c**, **f**) and quantitatively (**d**, **g**) decreases during days 1-5 of pluripotency exit after neuroectodermal (NE) differentiation trigger, closely mirroring the decrease of OCT4 protein, as assessed by immunofluorescence against endogenous OCT4 (‘anti-OCT4’). Cell samples were fixed at days 1, 2, 3, 4 or 5, as indicated. DAPI nuclear signal is shown as reference. In **d** and **g**, signals at days 1, 2, 3, 4, and 5 are normalized to that of non-triggered cells at day 0, and asterisks *** indicate statistically significant decrease compared to the day 0 values (Mann Whitney test p-value < 10^-3^, n > 6000 cells for each condition; boxplot whiskers indicate 99 percentile bounds). a.u.: arbitrary units. Scalebars: 50 μm. **e** and **h**, exoORACLE-OCT4 (**e**) and endoORACLE-OCT4 (**h**) levels correlate highly with OCT4 protein levels on a single-cell basis (Spearman’s correlation coefficient shown, n > 6000 cells for each condition).

### ORACLE-OCT4 reports OCT4 expression with high fidelity at single-cell level

We used both strategies to generate live exogenous ORACLE (‘exoORACLE’) and endogenous ORACLE (‘endoORACLE’) reporters of OCT4. OCT4 is a pluripotency TF widely used as a marker of pluripotency, whose expression is both necessary to establish and maintain the pluripotent state and lost when cells exit pluripotency ^40^. We tagged POM121 with fluorescent tdTomato and stably knocked-in exo and endo constructs in hESCs (see Methods). exoORACLE-OCT4 and endoORACLE-OCT4 reporters’ expression led to bright nuclear rim-localised signal in all cells, which after triggering neural differentiation was lost after 5 days, as would be expected for OCT4 (Fig. 1a and b (right) and Supplementary Fig. 1c and d).

To assess how faithfully ORACLE-OCT4 expression reports endogenous OCT4 expression, we induced neural differentiation and fixed cells at days 0 (untreated), 1, 2, 3, 4 and 5 following neural induction, and then analysed samples for ORACLE-OCT4 level and OCT4 immunofluorescence. We performed comparative quantitative analysis both at a cell population level and on a cell-by-cell basis for thousands of cells. As shown in Fig. 1c-h, the signals of both exoORACLE-OCT4 and endoORACLE-OCT4 gradually decreased throughout the time course, most visibly starting on day 3 of differentiation, similarly to OCT4 protein levels. This data indicate that ORACLE-OCT4 loss mirrors OCT4 loss at the population level (Fig. 1c-d, f-g). When we compared ORACLE-OCT4 signal and OCT4 protein level at a single-cell level, we found a high degree of correlation for both constructs (Fig. 1c and f). This correlation was particularly high for endoORACLE-OCT4, where it was typically >0.86 and up to 0.93. Thus, ORACLE levels quantitatively mirror transcription factor levels and changes at a single-cell level, indicating that they can be used to indirectly monitor TF levels with high fidelity.

### Visualizing spatiotemporal cell-to-cell heterogeneity during pluripotency exit in hPSCs

We then sought to investigate OCT4 spatiotemporal dynamics at the single-cell level by multi-day time-lapse microscopy. To this end, we generated an endoORACLE-OCT4 cell line co-expressing a histone H2B reporter tagged with the far red fluorescent reporter protein miRFP670 ^41^ (H2B-miRFP670) by CRISPR knock-in. This approach allowed us to simultaneously visualize chromatin and proliferation in addition to the OCT4-related signal through multiple cell cycles (Fig. 2a and Supplementary Video 1). Specifically, by imaging cells by time-lapse multi-colour confocal fluorescence microscopy for 5 days following neural induction, we monitored without interruption quantitative OCT4 level before, during, and after OCT4 disappearance in single-cells and their progeny during pluripotency exit. To minimize phototoxicity to cells we developed an imaging modality to differentially sample in time the two fluorescent reporters, by imaging the H2B-miRFP670 signal every 10 minutes and only imaging the ORACLE-OCT4 signal every 30 minutes. In this way we were able to grow the cells under the microscope with a proliferation rate indistinguishable from non-imaged cells (not shown).

**Figure 2.**
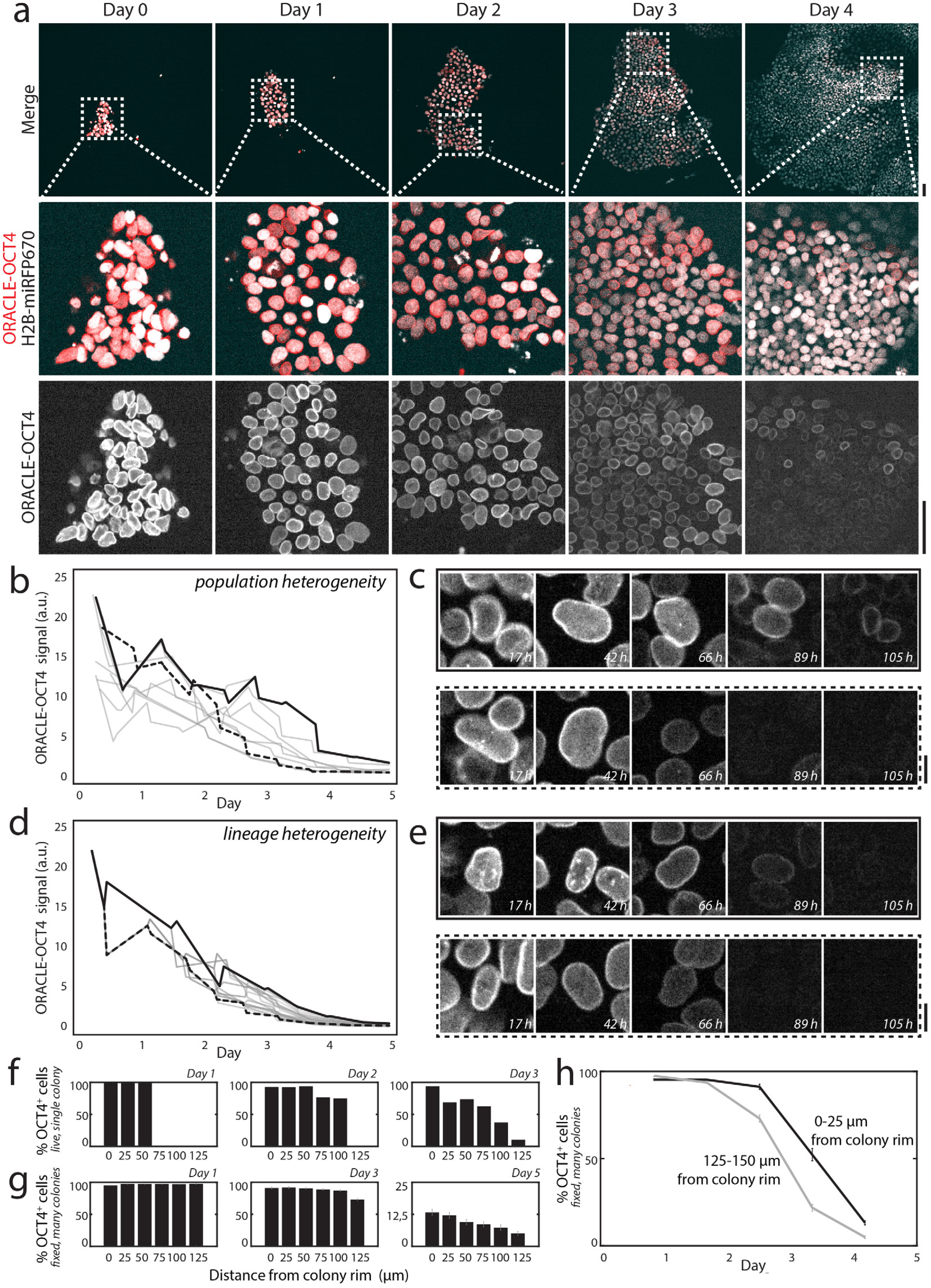
Spatiotemporal dynamics of ORACLE-OCT4 at single-cell level, by multi-day time-lapse microscopy. **a,** Image gallery of cells co-expressing ORACLE-OCT4 and the live chromatin reporter H2B-miRFP670 imaged continually by optimized, multi-day timelapse microscopy for 5 days after NE differentiation trigger at day 0. Dotted box and lines in the top row indicate areas magnified below. Scalebars: 50 μm. **b**, ORACLE-OCT4 signal intensity of 10 different cells in the population tracked through the 5 days of pluripotency exit exemplifying broad heterogeneity in the pattern of OCT4 loss among cells. The black dotted line shows the OCT4 loss pattern of a cell that loses most of its OCT4 before day 3, while the black solid line shows a cell that instead loses OCT4 only around day 4, some 24h later. **c**, Image gallery of the cells corresponding to the signals shown in **b** as a black dotted line (bottom) and a black solid line (top). **d**, ORACLE-OCT4 signal intensity of 177 cells originating from the same progenitor tracked through the 5 days of pluripotency exit, exemplifying heterogeneity in OCT4 loss pattern among cells stemming from the same lineage. The black dotted line shows a cell that loses most of its OCT4 by day 4, while the black solid line shows a cell that instead loses it only by day 5, some 24h later. **e**, Image gallery of the cells corresponding to the signals shown in **d** as a black dotted line (bottom) and a black solid line (top). **c** and **e**, time after differentiation trigger shown at the bottom of the image panels; Scalebars: 10 μm. **f**, **g** and **h**, Percentage of OCT4 positive (OCT4+) cells at different distances (‘0’: 0-25 μm, ‘25’: 25-50 μm, ‘50’: 50-75 μm, ‘75’: 75-100 μm, ‘100’: 100-125 μm, or ‘125’: 125-150 μm) from the colony rim during NE differentiation. **f** shows data taken from a time-lapse sequence of an individual colony at Days 1, 2 and 3. **g** shows collective time-course data taken from many colonies at Days 1, 3 and 5. **h** displays the percentage of OCT4+ cells at the colony rim (0-25 μm) versus away from the rim (125-150 μm) as a function of days of differentiation, based on the data of **g**. As can be seen from the plots, differentiating cells at the rim of colonies become OCT4-later than away from the rim, but with a similar kinetics of OCT4 loss.

Pluripotent cells have been shown to express heterogeneous levels of OCT4 before on-set of differentiation ^24^. Interestingly, through manual tracking of individual cells and their progeny throughout the 5 days of differentiation, we observed broad cell-to-cell heterogeneity also in the temporal pattern of OCT4 loss. Some differentiating hPSCs lost most of their OCT4 signal already after 2.5 days, while others only did so only after 4 days, i.e. more than 1 cell cycle later (Fig. 2b and c). Importantly, such heterogeneity could also be observed between progeny from the same lineage (Fig. 2d and e). This indicates that the cell of origin ^24^ alone cannot account wholly for the dynamics of pluripotency exit in hPSCs. Notably, we observed that as pluripotency exit proceeds, cells at the periphery of colonies appeared to retain higher OCT4 levels for longer than cells away from the periphery (see Fig. 1c, 1f and 2a, Day 3 and 4 panels). We confirmed this observation by quantitative fixed-cell time course and time-lapse analysis, which showed that peripheral cells lose OCT4 later than cells away from the colony rim (Fig. 2f-h and Supplementary Figs. 2a-d). Interestingly, however, despite this delay, once OCT4 levels began dropping, they did so with a similar kinetics independently of the cells’ location (Fig. 2h and Supplementary Figs. 2c and d).

Thus, spatial location is likely a key determinant of the timing, but not the kinetics, of pluripotency exit.

### Co-visualizing cell cycle changes and pluripotency exit dynamics at single-cell level

We then sought to exploit the ORACLE system to investigate how hPSCs coordinate cell cycle dynamics with pluripotency exit. To this end, we generated a triple CRISPR knock-in cell line co-expressing endoORACLE-OCT4, H2B-miRFP670 and the two-colour FUCCI cell cycle reporter ^29^ (Fig. 3a and Supplementary Video 2). This reporter combination made it possible to clearly distinguish the ORACLE-OCT4 nuclear rim-bound signal from the FUCCI nuclear signal (Fig. 3b), demonstrating that ORACLE effectively multiplexes the number of compatible reporters that can be observed and quantified simultaneously in live cells. With this combination, we quantitatively tracked cells and their progeny without interruption through 3 cell cycle transitions - G2/M (Fig. 3c and Supplementary Fig. 3b), M/G1 (Fig. 3d and Supplementary Fig. 3c) and G1 to S/G2 (Fig. 3e and Supplementary Fig. 3d) - and through 5 days of differentiation, while at the same time tracking pluripotency exit by ORACLE-OCT4 expression. As before, fluorescent signals were sampled at different time intervals to avoid phototoxicity: H2B-miRFP670 was imaged every 10 minutes, while ORACLE-OCT4 and FUCCI were imaged every 30 minutes.

**Figure 3.**
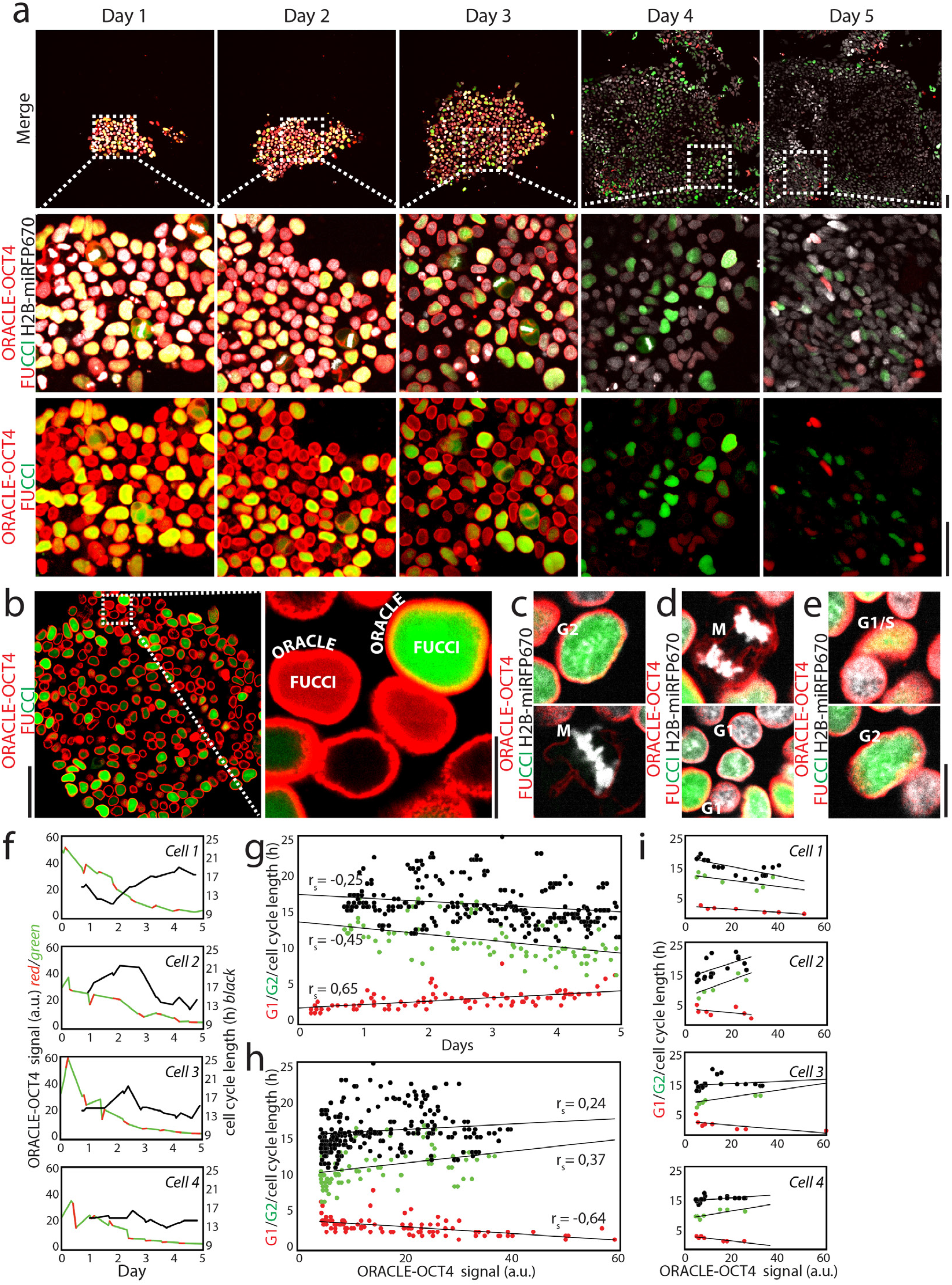
Co-visualization of cell cycle pattern and pluripotency exit dynamics at single-cell level. **a,** Image gallery of cells co-expressing ORACLE-OCT4, H2B-miRFP670 and the two-colour live FUCCI reporter of the cell cycle, imaged continually by optimized, multi-day time-lapse microscopy for 5 days following NE differentiation trigger at day 0. Dotted box and lines in the top row indicate areas magnified below. Scalebars: 50 μm. **b**, Cells co-expressing ORACLE-OCT4, H2B-miRFP670 and FUCCI, imaged at high (60x) magnification, showing that the nuclear rim-based ORACLE signal can be visually clearly distinguished from the nuclear FUCCI signal (only red and green fluorescence signals shown). Dotted box and lines on the left indicate the area magnified on the right image. Scalebars: 50 μm (left) and 10 μm (right). **c, d** and **e**, Consecutive time-lapse images of ORACLE-OCT4 H2B-miRFP670 FUCCI cells imaged at high (60x) magnification and undergoing the G2-M (**c**), M-G1 (**d**) and G1-S-G2 (**e**) transitions. **f**, Summary plots representing how OCT4 level (red/green line; left y axis) and cell cycle length (black line; right y axis) co-evolve through the 5 days following differentiation trigger, for 4 individual cells. OCT4 lines are coloured (red for G1, green for S/G2/M) to represent the progression of cell cycle phases as cells and their progeny progressively lose OCT4, based on the FUCCI readout. Cell cycle length is calculated as a running estimate, by measuring the time difference between a point in the cell cycle and the equivalent point at the next cell cycle. Note the heterogeneity in both OCT4 loss pattern and cell cycle pattern following differentiation trigger, among the 4 cells shown. **g** and **h**, G1 (‘G1; red dots, n=98), S/G2/M (‘G2’; green dots, n=96) and cell cycle (black dots, n=276) lengths as a function of days of differentiation (**g**) or OCT4 level (**h**), measured for 14 cells. Black lines: linear regressions; r_s_: Spearman rank-order correlation coefficient. **i**, G1 (red dots), S/G2/M (green dots) and cell cycle (black dots) lengths as a function of OCT4 level for the same 4 cells from **f**. Black lines: linear regressions.

We found that even at low magnification, the nuclear rim-bound red fluorescent ORACLE-OCT4 signal was clearly distinguishable from the FUCCI red fluorescence (Fig. 3a and Supplementary Fig. 3), making it straightforward to simultaneously quantify OCT4 level, S/G2 phase length (and consequently G1 phase length) and cell cycle duration on a singlecell basis through the 5 days of differentiation.

Similar to what we had observed with OCT4 variation, we found broad heterogeneity across individual cells in their cell cycle pattern following on-set of differentiation, with some cells shortening and others lengthening their cell cycle (Fig. 3f). In agreement with this data, cell cycle length across many cells correlated poorly with both time into differentiation (Fig. 3g; r_s_ = −0,25) and OCT4 level (Fig. 3h; r_s_ = 0,24). By contrast, G1 length was highly correlated with both time of differentiation and OCT4 levels (r_s_ = 0,65 and r_s_ = −0,64, respectively), whereby cells with highest OCT4 levels tended to have shortest G1 phase durations, in some cases as short as 1.5h or 2h. The correlation between OCT4 level and G1 length could be observed also at the single cell level (Fig. 3i). By contrast, neither cell cycle length nor S/G2 length correlated well with OCT4 level (Fig. 3i). Thus cell cycle length is a poor predictor of pluripotency exit. Conversely, G1 phase length is a good predictor of OCT4 level and therefore of pluripotency status during pluripotency exit, suggesting that these features are mechanistically linked.

### Multiplexed visualization of TF levels in live cells

Next, we set out to exemplify how ORACLE can be leveraged to enhance the number of different TF reporters that can be simultaneously visualized in cells. To this end, we generated a triple CRISPR knock-in cell line expressing endoORACLE-OCT4, H2B-miRFP670 and a live nuclear reporter of the pluripotency gene SOX2 ^42^. In the SOX2 construct, a mCherry-NLS protein is driven by the endogenous SOX2 promoter (pSOX2) by placing mCherry-NLS after the SOX2 coding sequence via the 2A system. As both the OCT4 (nuclear rim-localised) and SOX2 (nuclear localised) reporters are red fluorescent, this should allow testing whether two TF levels can be monitored simultaneously within a single fluorescence channel, effectively doubling the number of TF reporters that can be visualized in cells. Indeed, co-expression of pSOX2-driven mCherry-NLS with ORACLE-OCT4 confirmed that distinct OCT4 and SOX2 signals could be visualized within individual cells, despite being in the same fluorescence channel (Fig. 4a). Furthermore, quantitation of the SOX2 nuclear signal showed that it could be unequivocally distinguished from the nuclear background signal of cells expressing ORACLE-OCT4 alone (Fig. 4b). Given this specificity, we next examined the relative abundance of OCT4 and SOX2 with respect to each other on a cell-by-cell basis. By quantifying ORACLE-OCT4 and pSOX2-driven mCherry-NLS signals across a number of cells, we found that OCT4 and SOX2 levels are highly correlated (Fig. 4c), consistent with the known cooperative role and interaction between OCT4 and SOX2 in pluripotency control and in agreement with previous findings in mESCs ^23,43^. Taken together, these findings demonstrate that ORACLE allows for simultaneous observation and quantitation of dynamic behaviour of multiple TFs in live cells even within the same fluorescence channel, thereby multiplexing TF observation.

**Figure 4.**
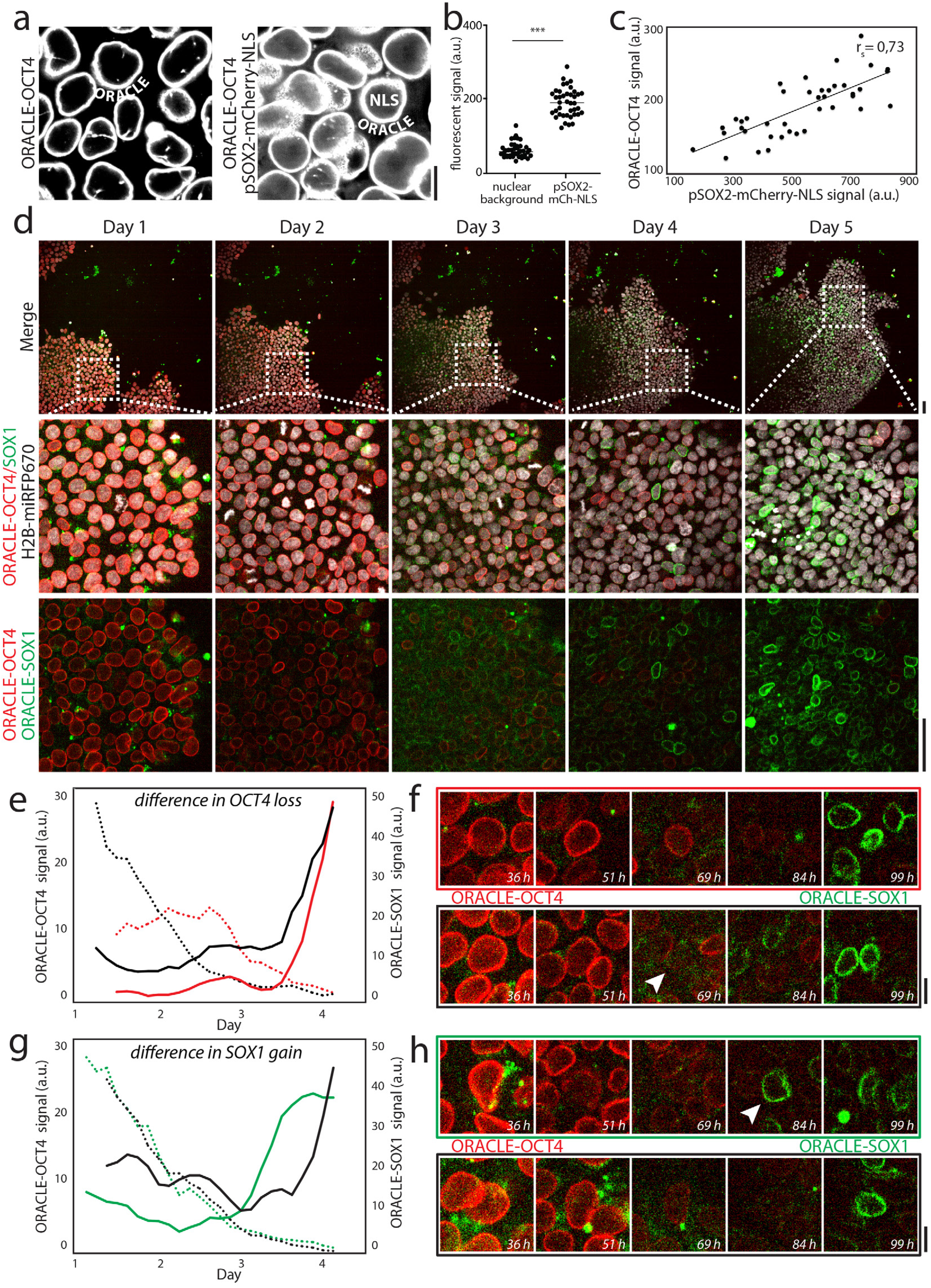
ORACLE multiplexes visualization of spatial and temporal TF dynamics in real time. **a**, Cells expressing ORACLE-OCT4 alone (left) or co-expressing ORACLE-OCT4 and the nuclear SOX2 reporter pSOX2-driven mCherry-NLS (right; labelled as ‘pSOX2-mCherry-NLS’), imaged at high (60x) magnification. As can be seen from the images, the nuclear rim-based ORACLE signal can be visually clearly distinguished from the nuclear mCherry-NLS signal (both signals correspond to red fluorescence, but are shown in grey for visual clarity). **b**, The SOX2 nuclear signal of cells co-expressing ORACLE-OCT4 and pSOX2-driven mCherry-NLS can be unequivocally distinguished from nuclear background signal of ORACLE-OCT4 expressing cells. asterisks *** indicate statistically significant difference (Kolmogorov Smirnov test p-value < 10^-3^, n=40 cells). **c**, OCT4 and SOX2 levels correlate in hPSCs. Black line: linear regression; rs: Spearman rank-order correlation coefficient. **d**, Image gallery of cells co-expressing ORACLE-OCT4, ORACLE-SOX1 and H2B-miRFP670, imaged continually by optimized, multi-day time-lapse microscopy for 5 days following NE differentiation trigger at day 0. Dotted box and lines in the top row indicate areas magnified below. Scalebars: 50 μm. **e**, ORACLE-OCT4 (dotted lines, left Y axis) and ORACLE-SOX1 (solid lines, right Y axis) signal intensities from 2 tracked cells (black and red lines) that have different OCT4 loss pattern (~20 hour difference) but similar SOX1 gain pattern. **f**, Image gallery of the cells corresponding to the signals shown in **e** (black/red boxes surrounding the images correspond to the coloured lines areas in **e**) illustrating how one cell (arrowhead) loses OCT4 earlier than the other. **g**, ORACLE-OCT4 (dotted lines, left Y axis) and ORACLE-SOX1 (solid lines, right Y axis) signal intensities from 2 tracked cells (black and green lines) that have similar OCT4 loss pattern but different SOX1 gain pattern (~20 hour difference). **h**, Image gallery of the cells corresponding to the signals shown in **g** (black/green boxes surrounding the images correspond to the coloured lines areas in **g**) illustrating how one cell (arrowhead) gains SOX1 earlier than the other. **f** and **h**, time after differentiation trigger shown at the bottom of the image panels; Scalebars: 10 μm.

### Visualizing live transcription factor ‘handover’ at single-cell level

Finally, we sought to exploit the ORACLE system to reveal the dynamics of transition between different cell states ‘live’ at single-cell level. To this end, we selected SOX1, a neuroectoderm TF and fate determinant ^44^, and designed an endoORACLE-SOX1 reporter. Using this reporter we generated a triple CRISPR knock-in cell line co-expressing endoORACLE-OCT4, endoORACLE-SOX1 and H2B-miRFP670 (Fig. 4d, and Supplementary Video 3). We then induced neural differentiation in these cells to visualize the transition from pluripotency to early neural stem cell fate in individual cells, i.e. to observe the dynamics of OCT4 loss and SOX1 gain, hereafter referred to as OCT4-to-SOX1 ‘handover’. Importantly, use of H2B-miRFP670 allowed us to track without interruption single-cells and their progeny even in instances where they had neither OCT4 nor SOX1 signal, allowing us e.g. to capture dividing cells even if they had lost OCT4 and compare the SOX1 expression patterns of their daughter cells. To minimize phototoxicity, fluorescent signals were again sampled at different intervals (H2B-miRFP670 imaged every 10 minutes, ORACLE-OCT4 and ORACLE-SOX1 imaged every 3 hours). Imaged colonies carrying this combination of reporters gradually lost OCT4 and gained SOX1 on differentiation, as expected (Fig. 4d).

Uninterrupted tracking of cells and their progeny throughout the 5 days of differentiation revealed that, just like for OCT4, cells display an ample range in the dynamics of SOX1 expression. Some cells switched on SOX1 already at day 3, while others did so at day 4 or even later (Fig. 4d). Unexpectedly, we found that the timing of OCT4 loss in cells occurred independently of the onset of SOX1 expression. Specifically, we observed instances where two cells had a similar timing of SOX1 activation but a different timing of OCT4 loss (Figs. 4e, f), and conversely, instances where cells with different timing of SOX1 activation had a similar pattern for OCT4 loss (Figs. 4g, h). This indicates that hPSCs display broad differences in how they coordinate the transition between pluripotency exit and onset of early differentiation. It also suggests that OCT4 and SOX1 are not mechanistically coupled in this fate transition. Thus, the ORACLE system is equally capable of detecting and can help distinguish between TF-driven mechanisms that are and that are not coupled.

## DISCUSSION

In this study, we expand the number of available ‘live’ fate and proliferation reporters that can be used to simultaneously and quantitatively monitor cellular behaviour. Our choice to localise ORACLE at the nuclear rim makes it particularly well suited for hPSCs, which grow as dense cell colonies with difficult to visualize compact cytoplasm. However, the ORACLE system can be easily modified in other cellular contexts to alternative (e.g. cytoplasmic, cell membrane) localisations. To exemplify the breadth of ORACLE’s multiplexing capacity, we combined it with H2B, FUCCI and nuclear TF probes. This shows that ORACLE can easily be combined with almost any type of reporter, including a wide variety of nuclear (e.g. signalling like ERK-KTR ^45^ or YAP/TAZ ^46^) and cytoplasmic (e.g. cytoskeletal, polarity and metabolic) cell biological reporters and biosensors ^47^. Because ORACLE exploits TF promoter-driven gene expression but does not involve tagging or mutation of TFs *per se*, it is not detrimental to cells ^22^ and does not modify pluripotency or differentiation propensity, and because it relies on targeted CRISPR knock-in it allows for control both of knock-in locus and copy number. This makes it an excellent non-invasive alternative to fluorescent TF fusion reporters. Furthermore, ORACLE’s modular design is adaptable to any TF or gene of interest. The sum of all these features makes ORACLE a flexible tool well suited both for basic discovery as well as tissue engineering and quality control applications ^48,49^.

The enhanced multiplexing capacity of ORACLE will enable studies of previously inaccessible regulatory layers, such as coordination of cell cycle/proliferation and cell fate transitions, TF handover and TF stoichiometry at single-cell level. To illustrate this we provide four examples that all warrant future investigation. In the first, we quantitated OCT4 levels during pluripotency exit and early NE differentiation by combining ORACLE-OCT4 with a H2B-miRFP670 reporter, allowing unequivocal tracking of cells before, during and after loss of the TF signal. We revealed broad cell-to-cell heterogeneity in OCT4 dynamics during early NE differentiation, in agreement with previously reported heterogeneity during early extraembryonic mesoderm differentiation ^24^. In addition, we showed that OCT4 loss does not happen uniformly within the differentiating tissue, and in particular happens later in cells in the periphery of cell colonies. This finding is in contrast with previous reports ^50^ and suggests that colony size and shape may have an unappreciated impact on differentiation efficiency and patterning ^51,52^. In the second example, we found that G1 phase length, and not cell cycle length, correlates with OCT4 level in hPSCs undergoing early NE differentiation. This finding indicates that cell cycle length *per se* is not informative about pluripotency exit status in hPSCs and reveals a tight connection between OCT4 and G1 length control that can now be investigated with the tools described here. This close relationship might be mediated via the CDK inhibitor (CKI) p21, as OCT4 has been shown to negatively regulate p21 in mouse embryonic stem cells ^18,53^, and/or by other mechanisms coupling OCT4 and the CDK4-6/CyclinD1-3 gearbox directly. Although OCT4 inheritance in daughter cells in hPSCs has been proposed to be a determinant for readiness to initiate differentiation (prior to presentation of any lineage cues) ^24^, our results suggest that this readiness may in fact stem from the underlying cell cycle properties and G1 length in cells. In a third example, we used an ORACLE-OCT4 pSOX2-driven mCherry-NLS system and confirmed that OCT4 and SOX2 abundances correlate at single-cell level, consistent with the known OCT4-SOX2 cooperativity in pluripotent stem cells. Finally, in a fourth example we used a dual ORACLE-OCT4/ORACLE-SOX1 system to probe directly TF handover at the single-cell level, an event that until now has never been visualized in real-time. With this system, we discovered that there is variable lag time between OCT4 loss and SOX1 gain, sometimes as long as 24h, which is within one typical cell cycle duration (equivalent to 1 generation) difference in time. These observations suggest that OCT4/SOX1-controlled early cell fate decisions are possibly more loosely coupled than previously appreciated, suggesting that cells exiting pluripotency may have a window of time before integrating or executing instructive fate signals. Taken together, these examples illustrate the power of this novel class of reporters in a wider range of applications.

Understanding how cells make complex and integrated cell fate decisions in space and time within tissues remains a quintessential topic in developmental biology and an important goal in regenerative medicine. Single-cell approaches, particularly scRNAseq and molecular barcoding technologies, have brought important advances in both areas, but as these are endpoint analysis-based, they are not able to reveal how cellular cause-effect decisions take place in real time in tissues. The tools and approach described here lend themselves naturally to large-scale ‘live’ microscopy phenotyping approaches aimed at generating comprehensive spatiotemporal maps of cell fate and proliferation dynamics in real time and space. We anticipate that such approaches will enable orthogonal avenues of research in stem cell biology and predictive synthetic tissue design.

## METHODS

### Maintenance and differentiation of hESCs

WA09 (H9) hESC line was purchased from WiCell (wicell.org) and maintained in Essential 8 medium (A151700, Thermo Fisher Scientific) on hESC-qualified growth factor reduced Geltrex-coated (A1413302, Thermo Fisher Scientific) 6 well plates. Cells were split into 6 well plates at 1:10 ratio when cells become confluent using 0.5 mM EDTA. Medium was changed everyday for maintenance of hESCs. To differentiate into neural stem cells (NSCs), essential 8 medium changed into PSC neural induction medium (A1647801, Thermo Fisher Scientific) on the next day after passage of hESCs. Cells were grown in the induction medium for 7 days to complete differentiation into NSCs then, the medium was changed every other day.

### Construction of plasmids for CRISPR/Cas9 mediated knock-in

Each pair of sgRNAs cloned into All-In-One (AIO) CRISPR/Cas9 nickase plasmids (#74119, Addgene) and sequences are shown in table 1. For the ORACLE-OCT4 (exo) construct, the fragments of 5’ and 3’ homology arms to target *AAVS1* locus were subcloned into hOCT4 promoter containing phOCT4-EGFP plasmid (#38776, Addgene) using AAVS1 SA-2A-puro-pA donor plasmid (#22075, Addgene) as a template. GFP fluorophore from the original plasmid was replaced with POM121 (p30554, Euroscarf) fused to tdTomato. ORACLE-OCT4 (endo) construct was modified from ORACLE-OCT4 (exo) as follows. The fragments of 5’ and 3’ homology arms of ORACLE-OCT4 (exo) were replaced using human OCT4-2a-eGFP-PGK-Puro (#31938, Addgene) to target endogenous *OCT4* locus with modification. Then, hOCT4 promoter was removed and POM121 fused tdTomato was placed before the stop codon of endogenous *OCT4* locus, followed by T2A to allow expression of target transcription factor genes together with nuclear rim expression. For the ORACLE-SOX1 construct, the fragments of 5’ and 3’ homology arms to target endogenous *SOX1* locus were synthesized as gene fragments (Eurofins genomics). Then, POM121 fused to 3 copies of GFP (p30474, Euroscarf) was placed before the stop codon of endogenous *SOX1* locus, followed by T2A sequence. SOX2-mCherry construct was modified from SOX2-t2a-mCherry (Plasmid #127538, Addgene) by replacing fragments of the 3’ homology arm to target the UTR region of SOX2 after stop codon using gene synthesis (Eurofins genomics) and inserting additional NLS sequences at the end of mCherry. For H2B-miRFP670 and FUCCI constructs, the fragments of 5’ and 3’ homology arms to target *ROSA26* locus were subcloned into H2B-670 (modified from pmiRFP670-N1, #79987, Addgene) or FUCCI (kind gift from Ludovic Vallier’s lab, U. of Cambridge). All of the cloning procedures were performed using In-fusion HD Cloning kit (639650, Takara) for seamless DNA cloning.

**Table 1.**
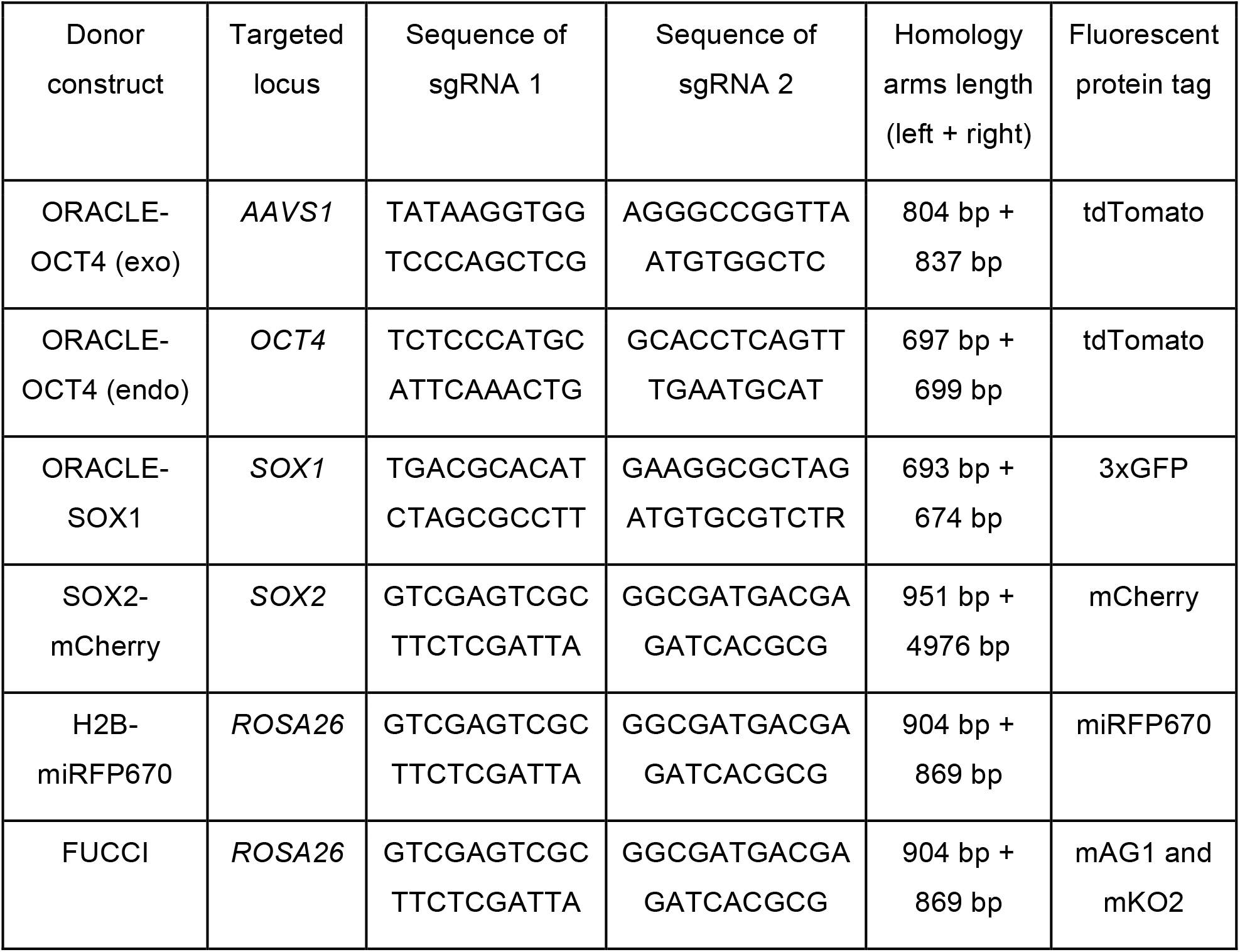
Specification of constructs for CRISPR/Cas9

### Generation of multi-reporter cell lines

To establish multi-colour reporter lines, cells were transfected with both AIO CRISPR/Cas9 nickase and donor vectors using lipofectamine stem transfection reagent (STEM00008, Thermo Fisher Scientific) according to the manufacturer’s protocol. Briefly, 1 μg of each plasmid, total 2 μg was diluted into 6 μl of reagent (1:3 ratio) in the 100 μl of Opti-MEM I medium (31985062, Thermo Fisher Scientific) and incubated for 15 min at RT. Transfected cells were dissociated with TrypLE™ Express Enzyme (12604013, Thermo Fisher Scientific) and re-suspended in BD FACS Pre-Sort Buffer (563503, BD biosciences) after 4 to 7 days of transfection. Live-cells were sorted on a BD Influx and collected into 1.5 ml microcentrifuge tubes containing DMEM supplemented with 50% of Knockout serum replacement (10828028, Thermo Fisher Scientific). Sorted cells were washed two times with 1 ml of Essential 8 medium and then, plated onto rhLaminin-521 (A29249, Thermo Fisher Scientific)-coated 96 well plates in the medium supplemented with 10 μM Y-27632 (SM02-1, Cambridge Bioscience) for 24 hr. Once cells were grown and established, they were subsequently transfected with additional constructs in a similar process to produce multicolour reporter cell lines.

### Immunofluorescence staining

Cells were fixed with 4% of paraformaldehyde (PFA) for 10 min and permeabilised with 0.3% Triton X-100 in phosphate buffer saline (PBS) for 15 min at room temperature. Cells were then blocked with 5% bovine serum albumin (BSA) in PBS for 2 hours, followed by an overnight incubation with primary antibodies to OCT4 (sc-5729, Santa Cruz Biotechnology) or SOX1 (4194, Cell Signaling Technology) at a 1:200 dilution in PBS containing 5% BSA at 4°C. Cells were then washed three times with 5% BSA PBS solution for 5 min each then, incubated with donkey anti-mouse or anti-rabbit conjugated to Alexa Fluor 405 or 488 secondary antibodies (Abcam) at a 1:500 dilution in PBS with 5% BSA) for 2 hours at room temperature in the dark.

### Time-lapse imaging

Established multi-colour hPSC lines were plated onto Geltrex-coated CellCarrier-96 Ultra Microplates (6055302, Perkin Elmer) a day before imaging supplemented with Essential 8 medium and maintained in the incubator. Differentiation trigger was applied by gently changing the medium into PSC neural induction just before placing the plate into the microscope. Cells were imaged using a Yokogawa CV7000 high throughput confocal microscope (Wako). For the experimental setting, ORACLE-OCT4 (endo) and SOX2-mCherry signal was captured every 30 min using 561 nm at 100 ms exposure time. ORACLE-SOX1 signal was captured every 3 hours using 488 nm at 500 ms. FUCCI signal was captured every 30 min using both 488 and 561 nm at 150 and 350 ms, respectively. Details are shown on table 2. Medium was changed every other day during the time-lapse of differentiation experiment for 5 days.

**Table 2.**
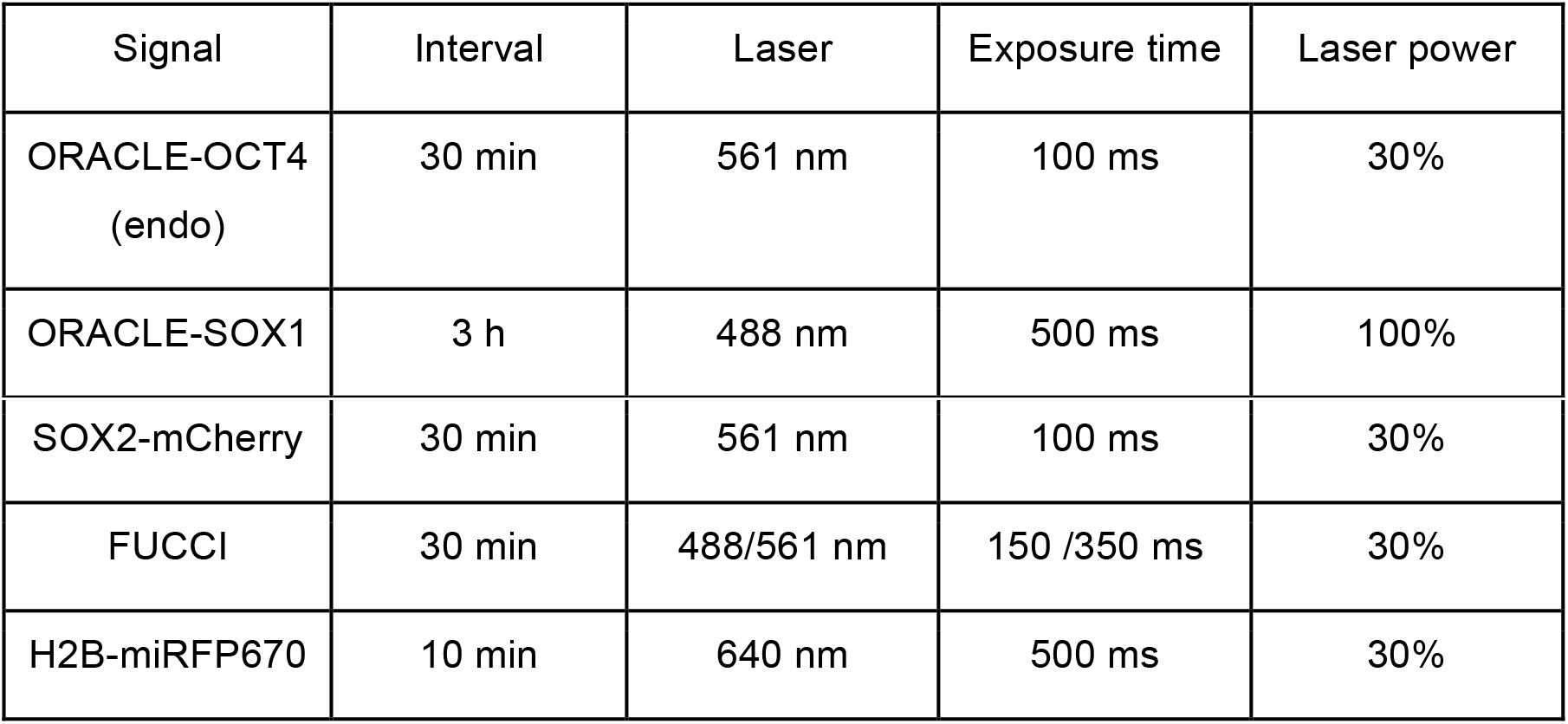
Experimental setting for time-lapse imaging

| Signal | Interval | Laser | Exposure time | Laser power |
| --- | --- | --- | --- | --- |
| ORACLE-OCT4<br>(endo) | 30 min | 561 nm | 100 ms | 30% |
| ORACLE-SOX1 | 3 h | 488 nm | 500 ms | 100% |
| SOX2-mCherry | 30 min | 561 nm | 100 ms | 30% |
| FUCCI | 30 min | 488/561 nm | 150 /350 ms | 30% |
| H2B-miRFP670 | 10 min | 640 nm | 500 ms | 30% |

### Quantitative image analysis

OCT4 protein, exoORACLE-OCT4 and endoORACLE-OCT4 signals quantitations from the immunofluorescence images shown in Figure 2 were carried out using a custom-made image processing pipeline using MATLAB (MathWorks). A median background value was calculated individually for each image and images were background subtracted, nuclei were segmented using the reference nuclear DNA DAPI signal, and the median fluorescence per pixel per nucleus was calculated for each nucleus for each channel (OCT4 protein, exoORACLE-OCT4 and endoORACLE-OCT4). Boxplot and bargraph statistical analyses were done in MATLAB, correlation analysis was done in R (https://www.r-project.org/). Uninterrupted cell tracking and signal quantitations for ORACLE-OCT4 (Figure 3), ORACLE-OCT4 and FUCCI (Figure 4), and ORACLE-OCT4 and ORACLE-SOX1 (Figure 5) were carried out by a combination of manual cell tracking and automated, background subtracted signal quantitations using custom scripts in FIJI (https://fiji.sc/).

## AUTHOR CONTRIBUTIONS

R.E.C.S. conceived the research and designed and supervised the study with help from S.K. and E.P. S.K. planned and performed reporter construction, vector construction, reporter knock-in and transgenic multi-reporter cell line generation as well as multi-day time-lapse microscopy experiments. E.R. did quantitative image and data analysis in MATLAB and R. R.E.C.S. and S.K. did manual cell tracking in FIJI and R.E.C.S. coded FIJI image analysis scripts. P.M.C. contributed to generating and modifying FUCCI constructs. R.E.C.S. wrote the paper with S.K and E.P.. Authors read and commented on the manuscript.

## ACKNOWLEDGEMENTS

We thank John Gurdon, Tony Kouzarides, Ludovic Vallier, Daniel Ortmann, Megan King, Sally Lowell, Guillaume Blin, and Dawei Sun for early discussions, Martin Hetzer, Jan Ellenberg, Nathalie Daigle, Ludovic Vallier, Sally Lowell, Guillaume Blin and Brian Burke for reagents, Marco Geymonat, Laura Wagstaff, Jonathan Lawson, Yasmin Paterson, Aubin Samacoits and Luc van Rensburg for early experimental/computational support or discussions, and Andrew Herman, Lorena Sueiro Ballesteros and the University of Bristol Flow Cytometry Facility for help with transgenic cell line generation. This work was supported by European Research Council Starting Researcher Investigator Grant SYSGRO (R.E.C.S.), University of Bristol funds (R.E.C.S., S.K.), Human Frontier Science Program (HFSP) grant RGP0043/2019 (R.E.C.S., S.K.), an EPSRC PhD studentship (E.R.), Gurdon Institute core funding from Wellcome Trust grant 092096 and Cancer Research UK grant C6946/A14492 (P.M.C.), Cancer Research UK Programme Foundation Award C38607/A26831 (E.P.) and Wellcome Trust Senior Research Fellowship 205010/Z/16/Z (E.P.).

## SUPPLEMENTARY INFORMATION

### SUPPLEMENTARY VIDEO CAPTIONS

**Supplementary Video 1**

5 day time-lapse video sequence of a hPSC cell line co-expressing endoORACLE-OCT4 and H2B-miRFP670, generated by CRISPR knock-in. To minimize phototoxicity to cells, an optimised imaging modality was used where H2B-miRFP670 signal was captured every 10 minutes and ORACLE-OCT4 signal every 30 minutes. This allowed cells to grow under the microscope with a proliferation rate indistinguishable from non-imaged cells, and enabled simultaneous visualization of chromatin, cell proliferation and OCT4 level dynamics through multiple cell cycles following neural induction at time 0.

**Supplementary Video 2**

5 day time-lapse video sequence of a hPSC cell line co-expressing endoORACLE-OCT4, H2B-miRFP670 and the two-colour FUCCI cell cycle reporter, generated by triple CRISPR knock-in. To minimize phototoxicity to cells, an optimised imaging modality was used where H2B-miRFP670 signal was captured every 10 minutes and ORACLE-OCT4 and FUCCI signals every 30 minutes. This allowed cells to grow under the microscope with a proliferation rate indistinguishable from non-imaged cells, and enabled simultaneous visualization of cell proliferation, OCT4 level dynamics and 3 cell cycle transitions - G2/M, M/G1 and G1 to S/G2 – through multiple cell cycles following neural induction at time 0.

**Supplementary Video 3**

5 day time-lapse video sequence of a hPSC cell line co-expressing endoORACLE-OCT4, endoORACLE-SOX1 and H2B-miRFP670, generated by triple CRISPR knock-in. To minimize phototoxicity to cells, an optimised imaging modality was used where H2B-miRFP670 signal was captured every 10 minutes and ORACLE-OCT4 and ORACLE-SOX1 signals every 3 hours. This allowed cells to grow under the microscope with a proliferation rate indistinguishable from non-imaged cells, and enabled visualization of the dynamics of OCT4-to-SOX1 ‘handover’ in individual cells, i.e. the loss of OCT4 followed by a gain of SOX1, indicative of the transition from the pluripotent state to the early neural stem cell fate live on a cell-by-basis.

**Supplementary Figure 1.**
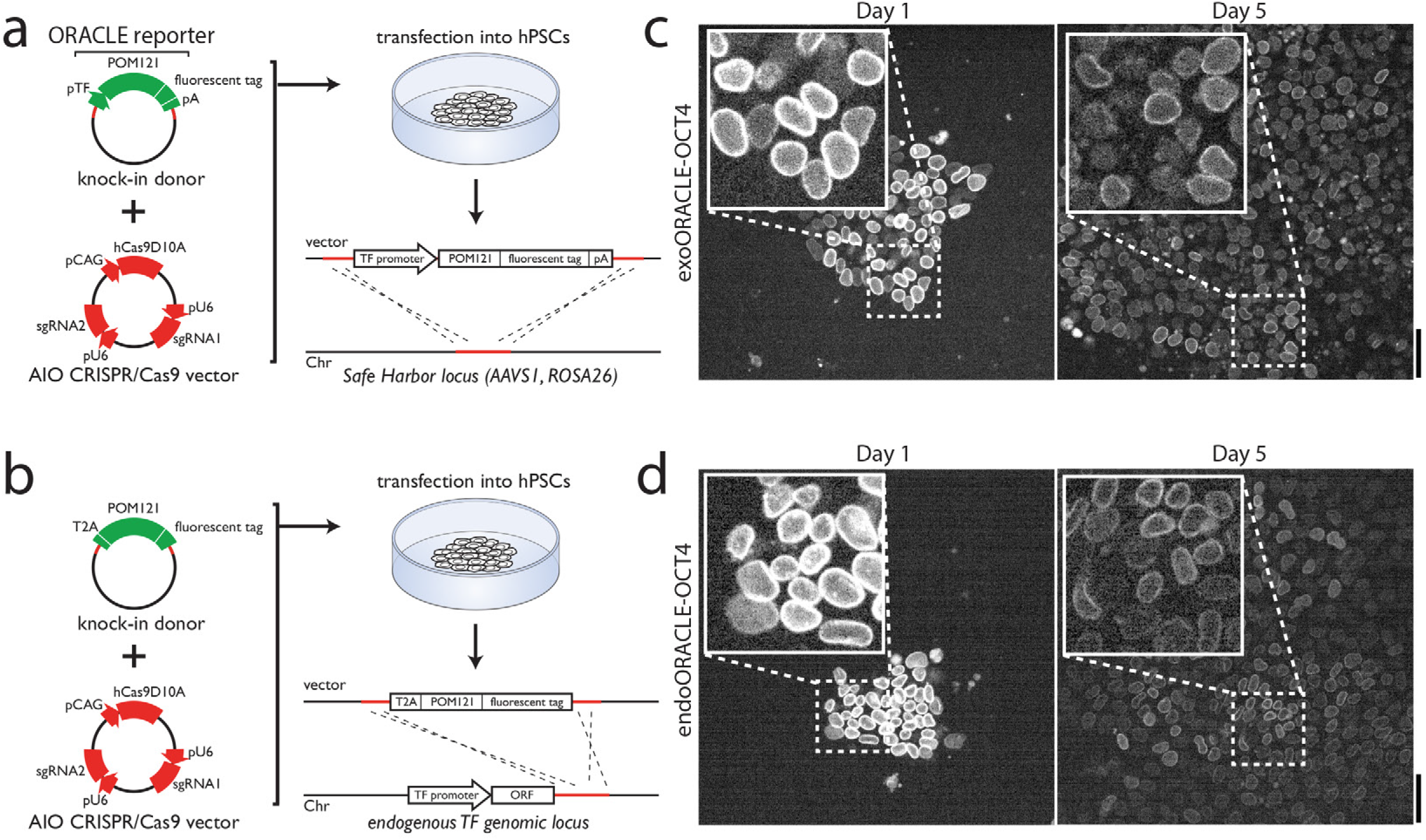
The ORACLE reporter system. **a,** Schematic of ‘exogenous’ ORACLE reporter design and knock-in in cells. A donor vector is generated that carries a construct containing the promoter sequence of a TF of choice (pTF), the POM121 gene fused to a fluorescent protein tagging sequence and a poly(A) signal (pA). The cassette is flanked at the 5’ and 3’ ends by homology sequences allowing targeting of the construct to a genomic safe harbour region (e.g. AAVS1 or Rosa26). Co-transfection of the donor vector with an All-In-One (AIO) CRISPR/Cas9 vector containing a sgRNA sequence(s) matching the homology region allows targeted, stable knock-in of the construct in cells, leading to pTF-driven expression from the exogenous safe harbour locus of fluorescent POM121 - this is the ‘exogenous’ ORACLE reporter (exoORACLE). **b,** Schematic of ‘endogenous’ ORACLE design and knock-in. A donor vector is generated that carries a construct containing a T2A sequence and the POM121 gene fused to a fluorescent protein tagging sequence. The cassette is flanked at 5’ and 3’ ends by homology sequences allowing targeting and tandem fusion of the construct to the 3’ end of a TF of choice after the end of the TF coding sequence. Co-transfection of the donor vector with an AIO CRISPR/Cas9 vector containing a sgRNA sequence(s) matching the TF 3’ homology region allows targeted, stable knock-in of the construct in cells, leading to expression of fluorescent POM121 driven by the endogenous TF’s promoter - this is the ‘endogenous’ ORACLE reporter (endoORACLE). **c,** and **d,** Images of hPSCs expressing exoORACLE-OCT4 (**c**) or endoORACLE-OCT4 (**d**), taken at days 1 and 5 of neuroectodermal (NE) differentiation. a.u.: arbitrary units. Dotted box and lines indicate area magnified in the insets. Scalebars: 50 μm.

**Supplementary Figure 2.**
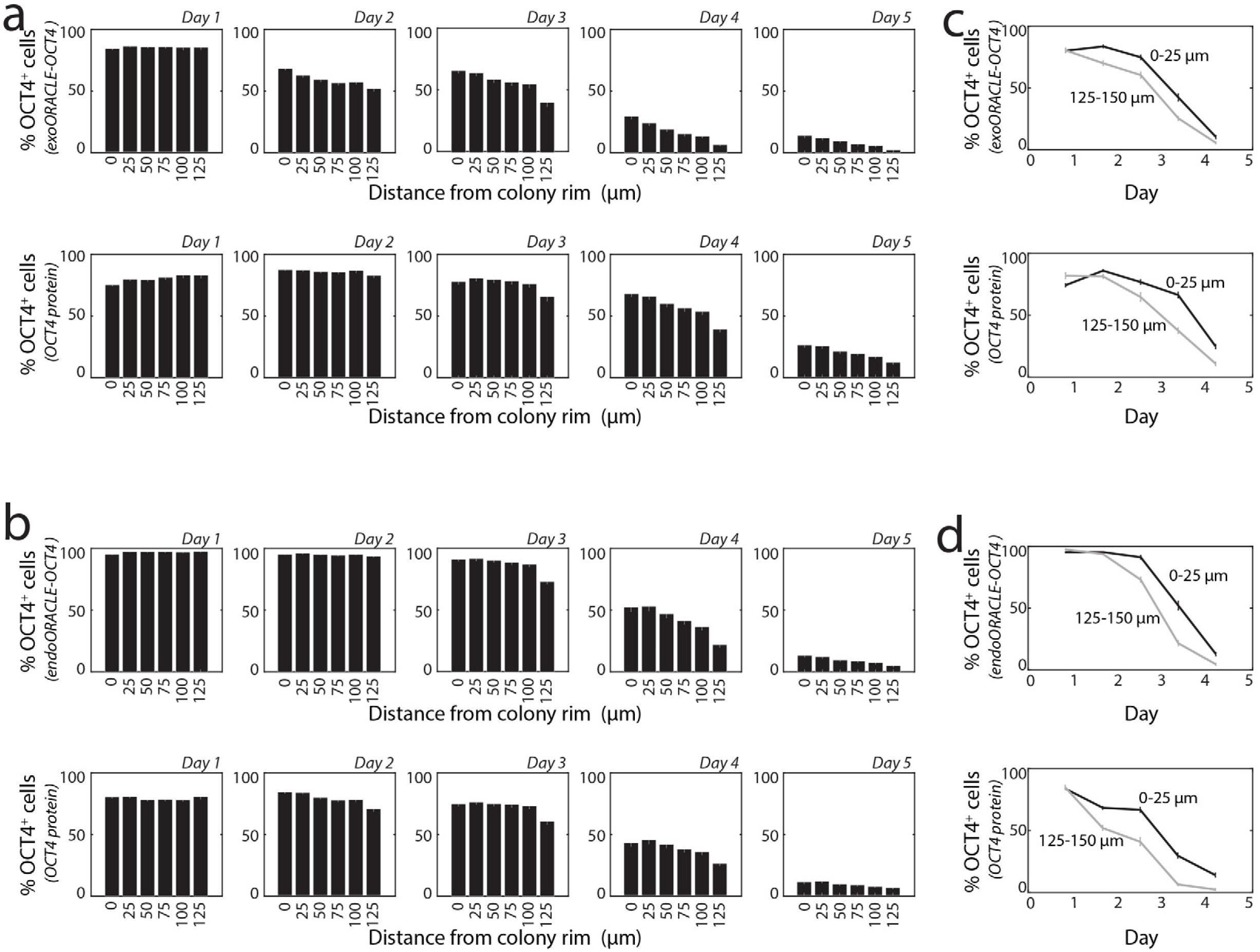
Differentiating cells at the rim of colonies lose OCT4 later than cells away from the colonies’ rim. **a** and **b,** Percentage of OCT4 positive (OCT4+) cells at different distances (‘0’: 0-25 μm, ‘25’: 25-50 μm, ‘50’: 50-75 μm, ‘75’: 75-100 μm, ‘100’: 100-125 μm, or ‘125’: 125-150 μm) from the colony rim during NE differentiation, taken from collective time-course data across many colonies. **a** shows data of exoORACLE-OCT4 expressing cells, fixed and processed by immunofluorescence against OCT4 protein. In the top row the percentage of OCT4+ cells is calculated based on the ORACLE-OCT4 signal; in the bottom row it is calculated based on OCT4 protein level. **b** shows data of endoORACLE-OCT4 expressing cells, fixed and processed by immunofluorescence against OCT4 protein. In the top row the percentage of OCT4+ cells is calculated based on the ORACLE-OCT4 signal; in the bottom row it is calculated based on OCT4 protein level. **c** and **d** display the percentage of OCT4+ cells at the colony rim (0-25 μm) versus away from the rim (125-150 μm) as a function of days of differentiation, based on the data from **a** and **b** respectively.

**Supplementary Figure 3.**
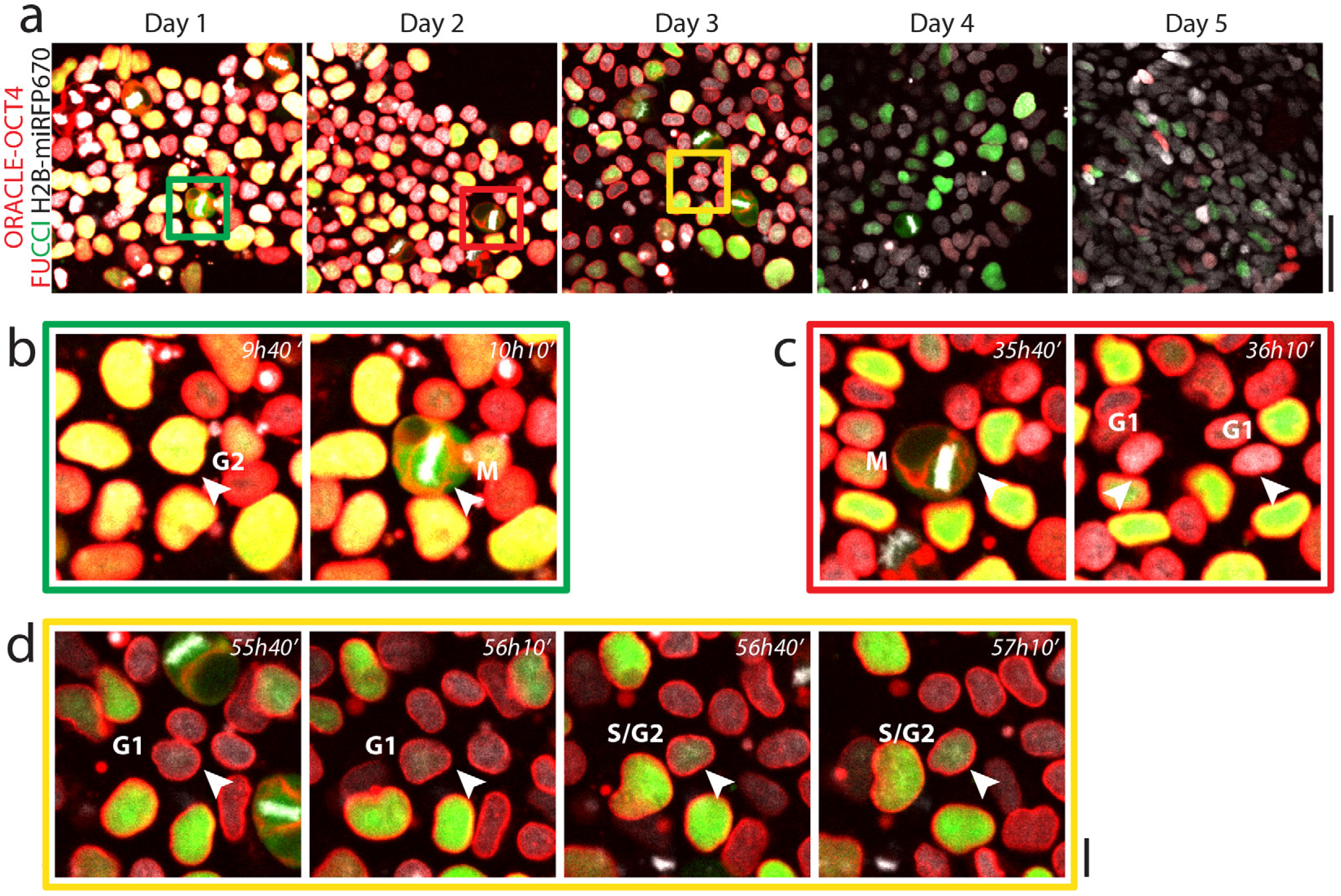
Co-visualization of cell cycle pattern and pluripotency exit dynamics at single-cell level. **a,** Image gallery from **Figure 3** of cells co-expressing ORACLE-OCT4, H2B-miRFP670 and the two-colour live FUCCI reporter of the cell cycle, imaged continually by optimized, multi-day time-lapse microscopy for 5 days following NE differentiation trigger at day 0. Coloured boxes indicate areas magnified below. Scalebar: 50 μm. **b**, **c** and **d**, Consecutive time-lapse images of ORACLE-OCT4 H2B-miRFP670 FUCCI cells undergoing the G2-M (**b**), M-G1 (**c**) and G1-S-G2 (**d**) transitions, for cells imaged at 20x magnification (cells indicated by arrowheads; green/red/yellow boxes surrounding the images correspond to the coloured boxed areas in **a**). Scalebar: 10 μm. Time after differentiation trigger shown at the top of the image panels. As can be seen from the images, the 3 transitions can be clearly seen at lower (20x) magnification.

## Notes

### Competing Interest Statement

The authors have declared no competing interest.

## REFERENCES

1 Tabar, V. & Studer, L. Pluripotent stem cells in regenerative medicine: challenges and recent progress. Nat Rev Genet 15, 82–92, doi:10.1038/nrg3563 (2014).

2 Goldfracht, I. et al. Generating ring-shaped engineered heart tissues from ventricular and atrial human pluripotent stem cell-derived cardiomyocytes. Nat Commun 11, 75, doi:10.1038/s41467-019-13868-x (2020).

3 Takebe, T. et al. Massive and Reproducible Production of Liver Buds Entirely from Human Pluripotent Stem Cells. Cell Rep 21, 2661–2670, doi:10.1016/j.celrep.2017.11.005 (2017).

4 Yiangou, L., Ross, A. D. B., Goh, K. J. & Vallier, L. Human Pluripotent Stem Cell-Derived Endoderm for Modeling Development and Clinical Applications. Cell Stem Cell 22, 485–499, doi:10.1016/j.stem.2018.03.016 (2018).

5 Zhao, C., Wang, Q. & Temple, S. Stem cell therapies for retinal diseases: recapitulating development to replace degenerated cells. Development 144, 1368–1381, doi:10.1242/dev.133108 (2017).

6 Cahan, P. & Daley, G. Q. Origins and implications of pluripotent stem cell variability and heterogeneity. Nat Rev Mol Cell Biol 14, 357–368, doi:10.1038/nrm3584 (2013).

7 Cuomo, A. S. E. et al. Single-cell RNA-sequencing of differentiating iPS cells reveals dynamic genetic effects on gene expression. Nat Commun 11, 810, doi:10.1038/s41467-020-14457-z (2020).

8 Friedman, C. E. et al. Single-Cell Transcriptomic Analysis of Cardiac Differentiation from Human PSCs Reveals HOPX-Dependent Cardiomyocyte Maturation. Cell Stem Cell 23, 586–598 e588, doi:10.1016/j.stem.2018.09.009 (2018).

9 Phillips, M. J. et al. A Novel Approach to Single Cell RNA-Sequence Analysis Facilitates In Silico Gene Reporting of Human Pluripotent Stem Cell-Derived Retinal Cell Types. Stem Cells 36, 313–324, doi:10.1002/stem.2755 (2018).

10 Breakthrough of the Year 2018., <http://vis.sciencemag.org/breakthrough2018/> (2018).

11 Torres-Padilla, M. E. & Chambers, I. Transcription factor heterogeneity in pluripotent stem cells: a stochastic advantage. Development 141, 2173–2181, doi:10.1242/dev.102624 (2014).

12 Chessel, A. & Carazo Salas, R. E. From observing to predicting single-cell structure and function with high-throughput/high-content microscopy. Essays Biochem 63, 197–208, doi:10.1042/EBC20180044 (2019).

13 Loeffler, D. & Schroeder, T. Understanding cell fate control by continuous single-cell quantification. Blood 133, 1406–1414, doi:10.1182/blood-2018-09-835397 (2019).

14 Neganova, I. et al. CDK1 plays an important role in the maintenance of pluripotency and genomic stability in human pluripotent stem cells. Cell Death Dis 5, e1508, doi:10.1038/cddis.2014.464 (2014).

15 Pauklin, S. & Vallier, L. The cell-cycle state of stem cells determines cell fate propensity. Cell 155, 135–147, doi:10.1016/j.cell.2013.08.031 (2013).

16 Gonzales, K. A. et al. Deterministic Restriction on Pluripotent State Dissolution by Cell-Cycle Pathways. Cell 162, 564–579, doi:10.1016/j.cell.2015.07.001 (2015).

17 Card, D. A. et al. Oct4/Sox2-regulated miR-302 targets cyclin D1 in human embryonic stem cells. Mol Cell Biol 28, 6426–6438, doi:10.1128/MCB.00359-08 (2008).

18 Lee, J., Go, Y., Kang, I., Han, Y. M. & Kim, J. Oct-4 controls cell-cycle progression of embryonic stem cells. Biochem J 426, 171–181, doi:10.1042/BJ20091439 (2010).

19 Goolam, M. et al. Heterogeneity in Oct4 and Sox2 Targets Biases Cell Fate in 4-Cell Mouse Embryos. Cell 165, 61–74, doi:10.1016/j.cell.2016.01.047 (2016).

20 Fowler, J. L., Ang, L. T. & Loh, K. M. A critical look: Challenges in differentiating human pluripotent stem cells into desired cell types and organoids. WIREs Dev Biol 9, doi:https://doi.org/10.1002/wdev.368 (2019).

21 Kirkeby, A. et al. Predictive Markers Guide Differentiation to Improve Graft Outcome in Clinical Translation of hESC-Based Therapy for Parkinson’s Disease. Cell Stem Cell 20, 135–148, doi:10.1016/j.stem.2016.09.004 (2017).

22 Etzrodt, M. & Schroeder, T. Illuminating stem cell transcription factor dynamics: longterm single-cell imaging of fluorescent protein fusions. Curr Opin Cell Biol 49, 77–83, doi:10.1016/j.ceb.2017.12.006 (2017).

23 Filipczyk, A. et al. Network plasticity of pluripotency transcription factors in embryonic stem cells. Nat Cell Biol 17, 1235–1246, doi:10.1038/ncb3237 (2015).

24 Wolff, S. C. et al. Inheritance of OCT4 predetermines fate choice in human embryonic stem cells. Mol Syst Biol 14, e8140, doi:10.15252/msb.20178140 (2018).

25 Strebinger, D. et al. Endogenous fluctuations of OCT4 and SOX2 bias pluripotent cell fate decisions. Mol Syst Biol 15, e9002, doi:10.15252/msb.20199002 (2019).

26 Piltti, K. M. et al. Live-cell time-lapse imaging and single-cell tracking of in vitro cultured neural stem cells - Tools for analyzing dynamics of cell cycle, migration, and lineage selection. Methods 133, 81–90, doi:10.1016/j.ymeth.2017.10.003 (2018).

27 Schroeder, T. Long-term single-cell imaging of mammalian stem cells. Nat Methods 8, S30–35, doi:10.1038/nmeth.1577 (2011).

28 Kimura, H. & Cook, P. R. Kinetics of core histones in living human cells: little exchange of H3 and H4 and some rapid exchange of H2B. J Cell Biol 153, 1341–1353, doi:10.1083/jcb.153.7.1341 (2001).

29 Sakaue-Sawano, A. et al. Visualizing spatiotemporal dynamics of multicellular cellcycle progression. Cell 132, 487–498, doi:10.1016/j.cell.2007.12.033 (2008).

30 Crisp, M. et al. Coupling of the nucleus and cytoplasm: role of the LINC complex. J Cell Biol 172, 41–53, doi:10.1083/jcb.200509124 (2006).

31 Wilhelmsen, K., Ketema, M., Truong, H. & Sonnenberg, A. KASH-domain proteins in nuclear migration, anchorage and other processes. J Cell Sci 119, 5021–5029, doi:10.1242/jcs.03295 (2006).

32 Gruenbaum, Y. & Foisner, R. Lamins: nuclear intermediate filament proteins with fundamental functions in nuclear mechanics and genome regulation. Annu Rev Biochem 84, 131–164, doi:10.1146/annurev-biochem-060614-034115 (2015).

33 Koch, A. J. & Holaska, J. M. Emerin in health and disease. Semin Cell Dev Biol 29, 95–106, doi:10.1016/j.semcdb.2013.12.008 (2014).

34 Daigle, N. et al. Nuclear pore complexes form immobile networks and have a very low turnover in live mammalian cells. J Cell Biol 154, 71–84, doi:10.1083/jcb.200101089 (2001).

35 Talamas, J. A. & Hetzer, M. W. POM121 and Sun1 play a role in early steps of interphase NPC assembly. J Cell Biol 194, 27–37, doi:10.1083/jcb.201012154 (2011).

36 Dultz, E. & Ellenberg, J. Live imaging of single nuclear pores reveals unique assembly kinetics and mechanism in interphase. J Cell Biol 191, 15–22, doi:10.1083/jcb.201007076 (2010).

37 Rabut, G., Doye, V. & Ellenberg, J. Mapping the dynamic organization of the nuclear pore complex inside single living cells. Nat Cell Biol 6, 1114–1121, doi:10.1038/ncb1184 (2004).

38 Ahier, A. & Jarriault, S. Simultaneous expression of multiple proteins under a single promoter in Caenorhabditis elegans via a versatile 2A-based toolkit. Genetics 196, 605–613, doi:10.1534/genetics.113.160846 (2014).

39 Daniels, R. W., Rossano, A. J., Macleod, G. T. & Ganetzky, B. Expression of multiple transgenes from a single construct using viral 2A peptides in Drosophila. PLoS One 9, e100637, doi:10.1371/journal.pone.0100637 (2014).

40 Wu, G. & Scholer, H. R. Role of Oct4 in the early embryo development. Cell Regen (Lond) 3, 7, doi:10.1186/2045-9769-3-7 (2014).

41 Shcherbakova, D. M. et al. Bright monomeric near-infrared fluorescent proteins as tags and biosensors for multiscale imaging. Nat Commun 7, 12405, doi:10.1038/ncomms12405 (2016).

42 Schaefer, T. & Lengerke, C. SOX2 protein biochemistry in stemness, reprogramming, and cancer: the PI3K/AKT/SOX2 axis and beyond. Oncogene 39, 278–292, doi:10.1038/s41388-019-0997-x (2020).

43 Thomson, M. et al. Pluripotency factors in embryonic stem cells regulate differentiation into germ layers. Cell 145, 875–889, doi:10.1016/j.cell.2011.05.017 (2011).

44 Abdelalim, E. M., Emara, M. M. & Kolatkar, P. R. The SOX transcription factors as key players in pluripotent stem cells. Stem Cells Dev 23, 2687–2699, doi:10.1089/scd.2014.0297 (2014).

45 Regot, S., Hughey, J. J., Bajar, B. T., Carrasco, S. & Covert, M. W. High-sensitivity measurements of multiple kinase activities in live single cells. Cell 157, 1724–1734, doi:10.1016/j.cell.2014.04.039 (2014).

46 Moya, I. M. & Halder, G. Hippo-YAP/TAZ signalling in organ regeneration and regenerative medicine. Nat Rev Mol Cell Biol 20, 211–226, doi:10.1038/s41580-018-0086-y (2019).

47 Science, A. I. f. C. Allen Institute for Cell Science Cell Catalogue <https://www.allencell.org/cell-catalog.html> (2020).

48 Chanthra, N. et al. A Novel Fluorescent Reporter System Identifies Laminin-511/521 as Potent Regulators of Cardiomyocyte Maturation. Sci Rep 10, 4249, doi:10.1038/s41598-020-61163-3 (2020).

49 Leigh, R. S., Ruskoaho, H. J. & Kaynak, B. L. A novel dual reporter embryonic stem cell line for toxicological assessment of teratogen-induced perturbation of anterior-posterior patterning of the heart. Arch Toxicol 94, 631–645, doi:10.1007/s00204-019-02632-1 (2020).

50 Narsinh, K. H. et al. Generation of adult human induced pluripotent stem cells using nonviral minicircle DNA vectors. Nat Protoc 6, 78–88, doi:10.1038/nprot.2010.173 (2011).

51 Blin, G. et al. Geometrical confinement controls the asymmetric patterning of brachyury in cultures of pluripotent cells. Development 145, doi:10.1242/dev.166025 (2018).

52 Laurent, J. et al. Convergence of microengineering and cellular self-organization towards functional tissue manufacturing. Nat Biomed Eng 1, 939–956, doi:10.1038/s41551-017-0166-x (2017).

53 Liu, L., Michowski, W., Kolodziejczyk, A. & Sicinski, P. The cell cycle in stem cell proliferation, pluripotency and differentiation. Nat Cell Biol 21, 1060–1067, doi:10.1038/s41556-019-0384-4 (2019).

